# “Wissenschaft fürs Wohnzimmer” – two years of interactive, scientific livestreams weekly on YouTube

**DOI:** 10.1101/2022.09.30.509854

**Authors:** Nicolas Stoll, Matthias Wietz, Stephan Juricke, Franziska Pausch, Corina Peter, Jana C. Massing, Miriam Seifert, Moritz Zeising, Melissa Käß, Rebecca McPherson, Björn Suckow

## Abstract

Science communication is becoming increasingly important to connect academia and society, and to counteract fake news among climate change deniers. Online video platforms, such as YouTube, offer great potential for low-threshold communication of scientific knowledge to the general public. In April 2020 a diverse group of researchers from the Alfred Wegener Institute Helmholtz Centre for Polar and Marine Research launched the YouTube channel “Wissenschaft fürs Wohnzimmer” (translated to “Sitting Room Science”) to stream scientific talks about climate change and biodiversity every Thursday evening. Here we report on the numbers and diversity of content, viewers, and presenters from 2 years and 100 episodes of weekly livestreams. Presented topics encompass all areas of polar research, social issues related to climate change, and new technologies to deal with the changing world and climate ahead. We show that constant engagement by a group of co-hosts, and presenters from all topics, career stages, and genders enable a continuous growth of views and subscriptions, i.e. impact. After 783 days the channel gained 30,251 views and 828 subscribers and hosted well-known scientists while enabling especially early career researchers to improve their outreach and media skills. We show that interactive and science-related videos, both live and on-demand, within a pleasant atmosphere, can be produced voluntarily while maintaining high quality. We further discuss challenges and possible improvements for the future. Our experiences may help other researchers to conduct meaningful scientific outreach and to push borders of existing formats with the overall aim of developing a better understanding of climate change and our planet.

## 1 Introduction

Communicating science in times of human-made climate change (IPCC, 2013, 2021), biodiversity loss (IPBES, 2019; Vaidyanathan, 2021), and the anticipated reaching of tipping points (Lenton et al., 2008, 2019) has become important for researchers and society. Global science-driven movements such as Fridays4Future (Hagedorn et al., 2019) and Scientists4Future underline the interest in and importance of well-communicated science. The global COVID-19 pandemic with its manifold implications for society and the link between science denial, e.g., in the form of climate change denial, and conspiracy thinking (e.g., Lahrach and Furnham, 2017; Uscinski et al., 2017) emphasises the need to communicate scientific results in an approachable way. Scientific knowledge gain is essential for politicians, economics, and society to take fact-based and appropriate decisions, especially during crises such as the COVID-19 pandemic and climate change.

Commonly, the gap between science and society has been bridged by the media rather than academia, often resulting in inaccurate or sensationalised results (Ladle et al., 2005). However, there are plenty of possibilities for scientists to reach an audience outside of academia and interact directly with the public. In addition to publishing in scholarly journals and presenting at conferences, alternative options such as blogs, homemade videos and the variety of social media activities enable a change in communication (Bickford et al., 2012) which makes research accessible and understandable to the public thereby directly linking science and society. Effective science communication allows people to make deliberate decisions by informing themselves about the risks and benefits of their actions. Instead of following the mantra “the public can’t understand science”, it should be “the public has little chance to learn science” (Fischhoff, 2013). In this study, Fischhoff presented the following aspects to determine if communication is efficient: it has to 1) contain the needed information, 2) exist in places the audience can access, and 3) offer an understandable format. In return, scientists can gain the public’s support by presenting the merit and trustworthiness of science (Fischhoff and Scheufele, 2013). Online science communication is a promising way to reach those goals, allowing to link publications or creating dialogues through comments and live discussions in forums or blogs, thus merging the real world with online interaction (Bubela et al., 2009). However, the main challenge remains to reach the audience and hence generate an impact.

Producing science communication content for established media platforms, e.g. in form of videos and podcasts, substantially facilitate reaching a broad audience. One of the most popular websites is YouTube, with more than one billion monthly users (Similarweb, 2022). It offers a place to upload commercial and homemade content covering sports, news, documentaries, music videos, tutorials and many more. This amount of information forces creators to understand how to capture the audience’s (long-term) attention and what factors contribute to success, i.e. channel popularity, views, continuous growth in subscribers, comments, ratings, and number of shares (Burgess et al., 2009). Factors impacting success are video length, continuity of content, reoccurring channel hosts, the pace of delivery, and whether the channel provides professionally- or user-generated content (Welbourne and Grant, 2016). Academic content is still only sporadically produced for platforms such as YouTube, despite their wider reach and hence large potential to communicate science (Maynard, 2021). The global spread of COVID-19 forced scientists to rethink established approaches of reaching the public, such as public presentations and discussion rounds, and make more use of online platforms. Thus, new challenges surfaced, such as 1) reaching an audience while most people were isolated at home, 2) developing concepts to attract people to attend online events, and 3) keeping this attention throughout months of pandemic-induced ups and downs.

In summer 2019 scientists from the Alfred Wegener Institute Helmholtz Centre for Polar- and Marine Research (AWI) formed AWIs4Future, a regional group within Scientists4Future. In April 2020, a team of 11 early career scientists (ECRs) from diverse disciplines active in AWIs4Future adapted the successful concept of the event series *Science goes Public* in the German towns Bremen and Bremerhaven to the internet. *Science goes Public* enables scientists to casually talk about their research in pubs and bars twice a year. These free-of-charge events attracted a wide audience, and were often fully booked. The pandemic forced *Science goes Public* to cancel all events. Thus, virtual alternatives had to be developed to continue science communication while public life came to an almost standstill.

“Wissenschaft fürs Wohnzimmer” (WfW), translated to “Sitting Room Science”, aimed to bridge this gap by hosting moderated, weekly YouTube livestreams focusing mostly on climate- and environmental-science-related topics. The main goals were: present current science topics of broad relevance, a low threshold for accessing the medium, free of charge for the audience, having a live and on-demand option, being interactive, and offering a relaxed, inclusive atmosphere. The skyrocketing use of video conferences across science demonstrates the potential for further use in areas such as science communication. Zoom as meeting software that offers connection with YouTube allowed hosting live presentations and direct interaction via the YouTube chat with viewers from around the world.

In this article we demonstrate the potential of establishing a low-cost, interactive, and accessible medium to communicate science to a broad audience. By sharing our experiences and best practices, we further aim to encourage other scientists to reach out to the public, contributing towards enhancing the quality of science communication, both between public and science 70 and among the scientific community.

## 2 Methods

### 2.1 Streaming via YouTube

The first stream took place on 16.04.2020 on the YouTube channel “*TRR 181 Energy transfers in Atmosphere and Ocean*” before the creation of the dedicated channel for WfW on 18.04.2020, roughly one month after the first strict lockdown in Germany. The original target audience was the science-interested public in the north of Germany (owing the location of AWI). Hence, the majority of talks are in German. In irregular intervals, WfW features presentations in English (12 to date). The presentations are generally between 20-45 min in length followed by 15 min of questions, aiming at a total streaming time of 45-60 min. Questions can be asked by the entire audience via live chat, only a free YouTube account is necessary. To enable additional interaction the audience is encouraged to add 80 comments afterwards, or contact WfW via the AWIs4Future accounts on Instagram and Twitter.

### 2.2 Live discussion via Zoom

A Zoom Pro account is used for a video call between three WfW hosts and one or two presenters. We choose Zoom Pro because of unlimited meeting time, direct streaming to YouTube, control over key settings (e.g., privacy), and communication among the host team via the chat function. However, discussions about using a non-commercial, open-source, and free broadcast alternative, such as OBS, are ongoing. The host team is usually gender-balanced consisting of scientists from different research disciplines and from different career stages ranging from new doctoral researchers to established PostDocs and senior scientists, mainly from AWI. The team meets 20-30 minutes prior to the livestream to perform a technical check and to discuss relevant details, such as the introduction, order of appearance and the way to announce next week’s presenter. Around 30s before the beginning of the livestream all conversations are ended and all microphones are muted. The stream is started, appearing with 30s delay to the live viewers. After playing the *WfW jingle*^1^, recorded by a WfW team member and band, one co-host welcomes the audience, introduces themselves, and the presenter. The other two co-hosts present themselves, and the presenter takes over.

During the presentation the presenter leads through their slides and is the only one speaking. Questions are normally answered after the presentation, based on the YouTube live chat monitored and moderated by one co-host. Occasionally, one of the co-hosts asks questions from the YouTube chat during the presentation, which can create a more lively and communicative atmosphere instead of a “lecture”. Depending on the audience the discussion can include a dozen questions ranging from basic to very specific. Spam and abusive comments occur rarely and are deleted and respective composers blocked as quickly as possible. The discussion usually ends after a maximum of 20 minutes and the WfW jingle concludes the livestream and the recording. In a debriefing the presenter is informed about the statistics, such as the number of live viewers and activity in the chat. A pleasant and often more in-depth discussion usually continues between the co-hosts and the presenter before the Zoom meeting is concluded.

### 2.3 Post-processing and on-demand

The recorded stream is automatically uploaded to YouTube. A few videos were edited slightly after the livestream, e.g. to cut out corrupted parts due to failed internet connection. The comment section below the videos remains open and is regularly monitored by the WfW team. Submitted questions are answered or, if necessary, forwarded to the presenter. Talks are sorted by content and placed in at least one of 9 different playlists (Table 1) for quick browsing.

**Table 1.** WfW playlists as of 09.06.22. Some videos appear in more than one playlist.

| Playlist | # videos |
| --- | --- |
| Klimawandel & Gesellschaft (*Climate Change & Society*) | 32 |
| Arktis & Antarktis (*Arctic & Antarctica*) | 26 |
| Technologien & Innovationen (*Technology & Innovation*) | 17 |
| Artenvielfalt (*Biodiversity*) | 16 |
| Englische Episoden (*English Episodes*) | 12 |
| Ernährung der Zukunft (*Diet of the Future*) | 10 |
| Modellierung von Ozean und Klima (*Modelling of Ocean and Climate*) | 10 |
| Permafrost und (Meeres)Boden (*Permafrost and (Sea)floor*) | 9 |
| MOSAiC (*MOSAiC*) | 2 |

## 3 Results

The WfW channel was created on 18.04.2020 with the first video originally streamed on the 16.04.2020 via a different channel (re-uploaded on the 19.04.2020). As of June 9 2022, WfW hosted 100 scientific livestreams (Fig. 1) whose stats are presented in this study. Streams took place every Thursday, except during Christmas, Easter, and during a one-month summer break in 2021.

**Figure 1.**
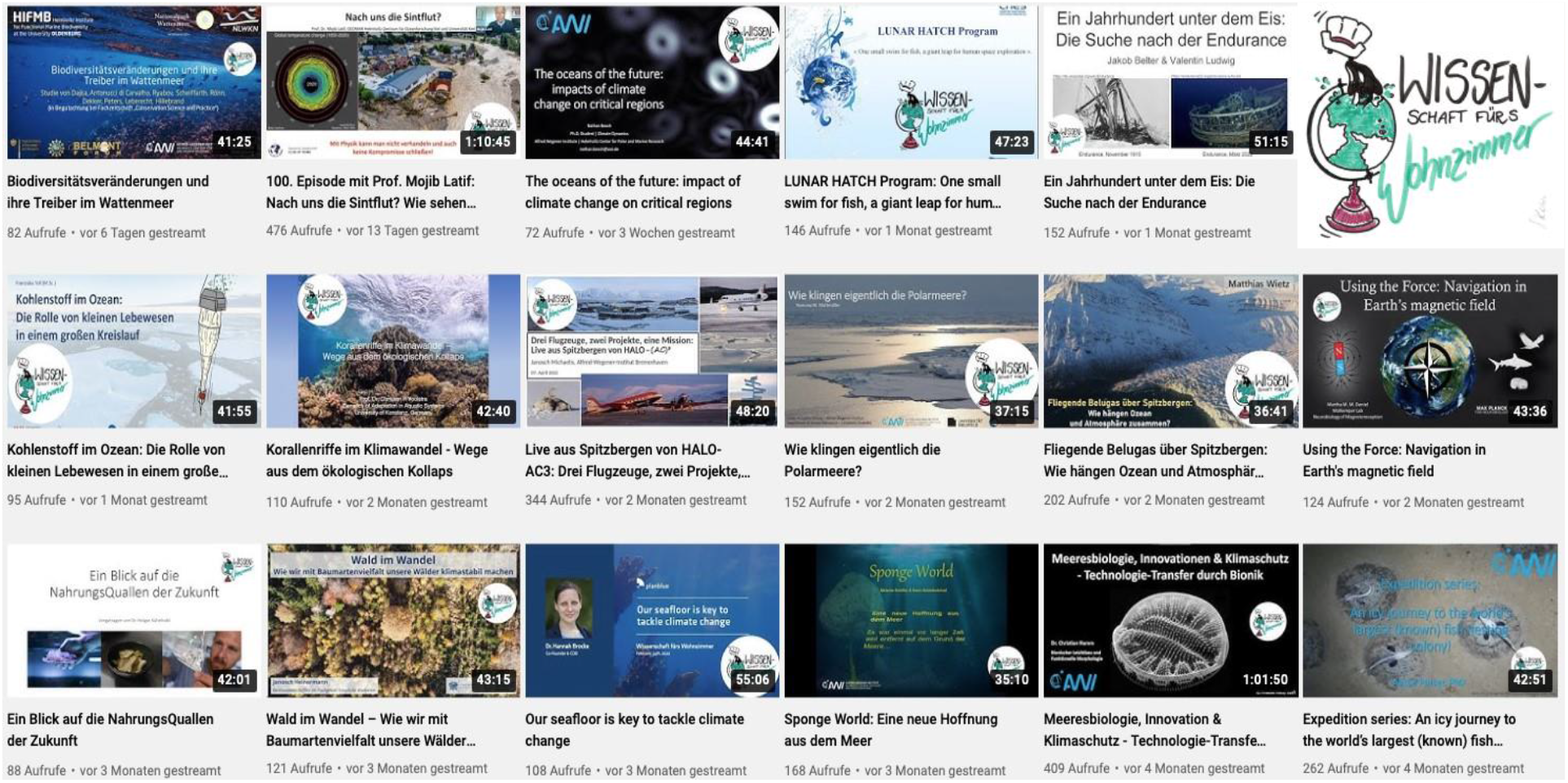
Overview of recently uploaded WfW videos as of 16.09.2022. The logo of the channel is shown in the upper right corner (credit: Svenja Kröner).

### 3.1 Views

In the analysed time frame (18.04.2020-09.06.2022) videos of the WfW channel gathered an accumulated number of 30,251 views (Fig. 2). Based on available YouTube information, one legitimate view is defined as intentionally clicking a video and watching a minimum of 30 s. There was a total of 10,645 views in 2020, 13,512 views in 2021, and 6,094 views until 09.06.2022 with a stable increase in cumulative views over the 783 days analysed (Fig. 2).

**Figure 2.**
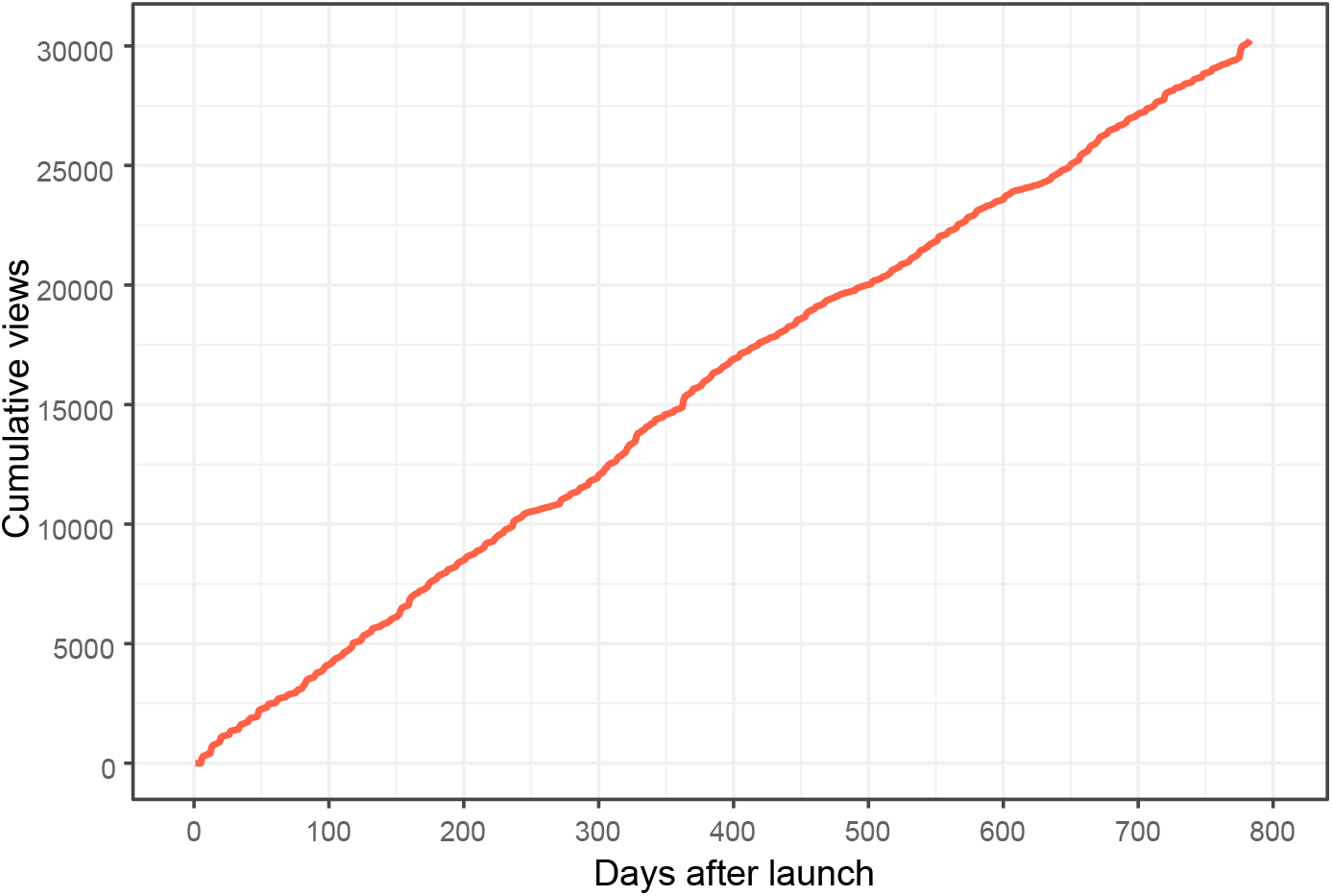
Cumulative views since the launch of the channel on 18.04.2020.

Daily views of the channel are up to 327 (Fig. 3). The lowest numbers were recorded in the Christmas, New Years, and summer breaks of 2020 and 2021. The highest numbers (326 and 327) were reached on the 50th and 100th episodes of WfW featuring Dr. Gregor Hagedorn (founder of Scientists for Future), and Prof. Mojib Latif (GEOMAR Helmholtz Centre for Ocean Research Kiel and University Kiel), respectively. The average and median number of views per day is 38.6 and 27, respectively (Fig. 3). In general, views are highest on Thursdays when new content is streamed live. Videos were watched for a total of 4,830 h. The cumulative view time per video ranges from 10.1 h to 146.2 h with an average play time of 5 min to 20 min.

**Figure 3.**
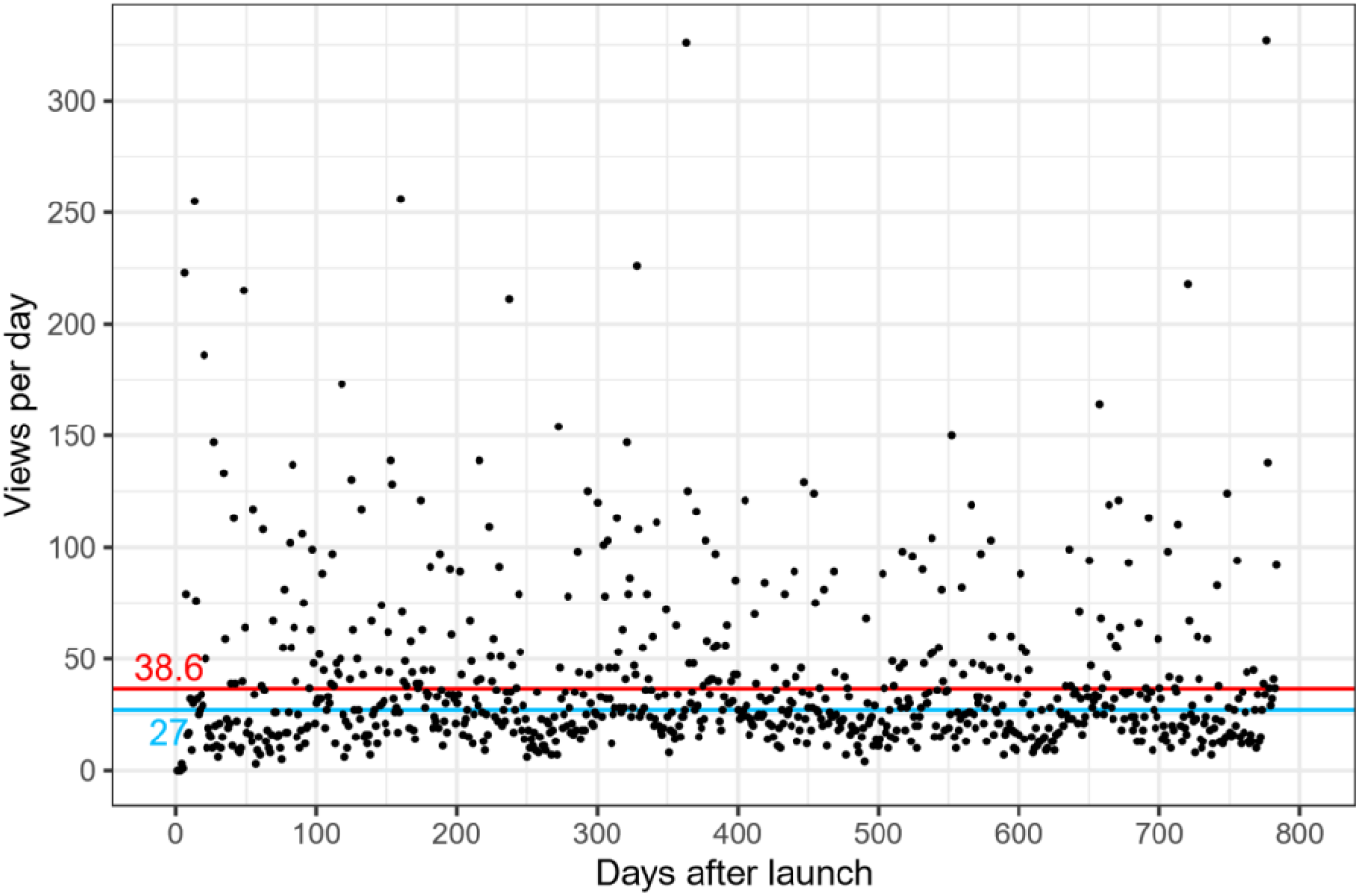
Views of the channel per day since the launch on 18.04.2020. The red line is the average number of views per day (38.7), the blue line is the median (27).

### 3.2 Subscriptions

WfW has 828 subscribers as of June 9, 2022 (Fig. 4B) corresponding to 0.9 subscriptions per day, 7.4 subscriptions per week, and 32.2 subscriptions per month. Subscriptions per day range from −3 to 26 (Fig. 4A). The strong increases in subscriptions (Fig. 4) coincide with presentations by the publicly known scientists Mojib Latif, Peter Lemke, and Gregor Hagedorn. Times of stagnation coincide with periods of channel inactivity, such as the summer breaks, Christmas and New Years 2020 and 2021. 74.6% of all views come from users without a subscription to WfW (22,573), 25.4% of the views are from WfW subscribers (7,678). Similarly, the total view time of non-subscribers is 3,364 h, whereas subscribers watched for a total time of 1,466 h (Fig. 5). Subscribers watch on average longer (around 11 min) than non-subscribers (around 9 min) (Fig. 5). Most subscribers were gained in connection with the presentations of Gregor Hagededorn and Mojib Latif, with 19 and 18 new subscriptions per video, respectively.

**Figure 4.**
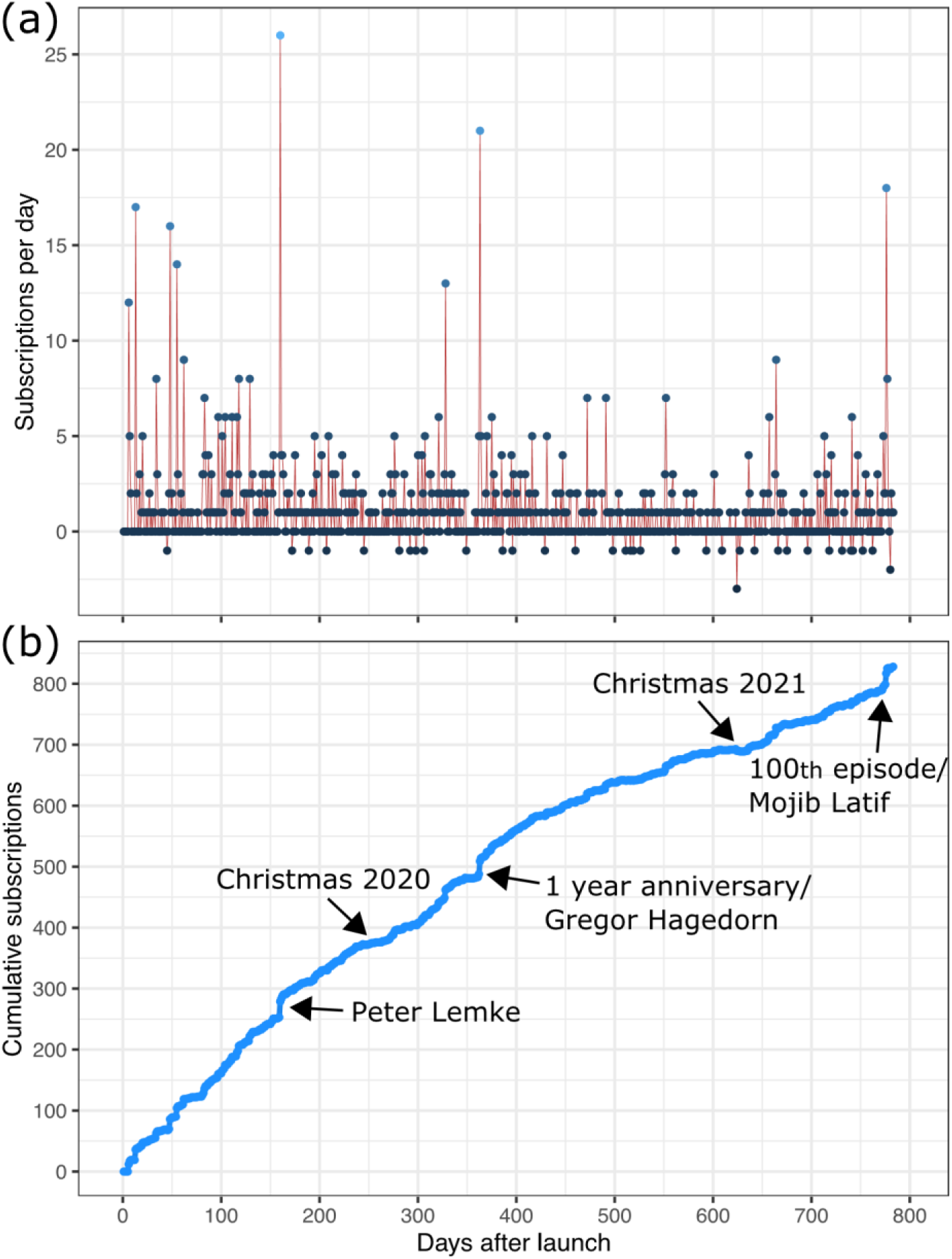
A) Subscriptions gained/lost per day. B) Cumulative amount of subscriptions. Marked are high and low impact events.

**Figure 5.**
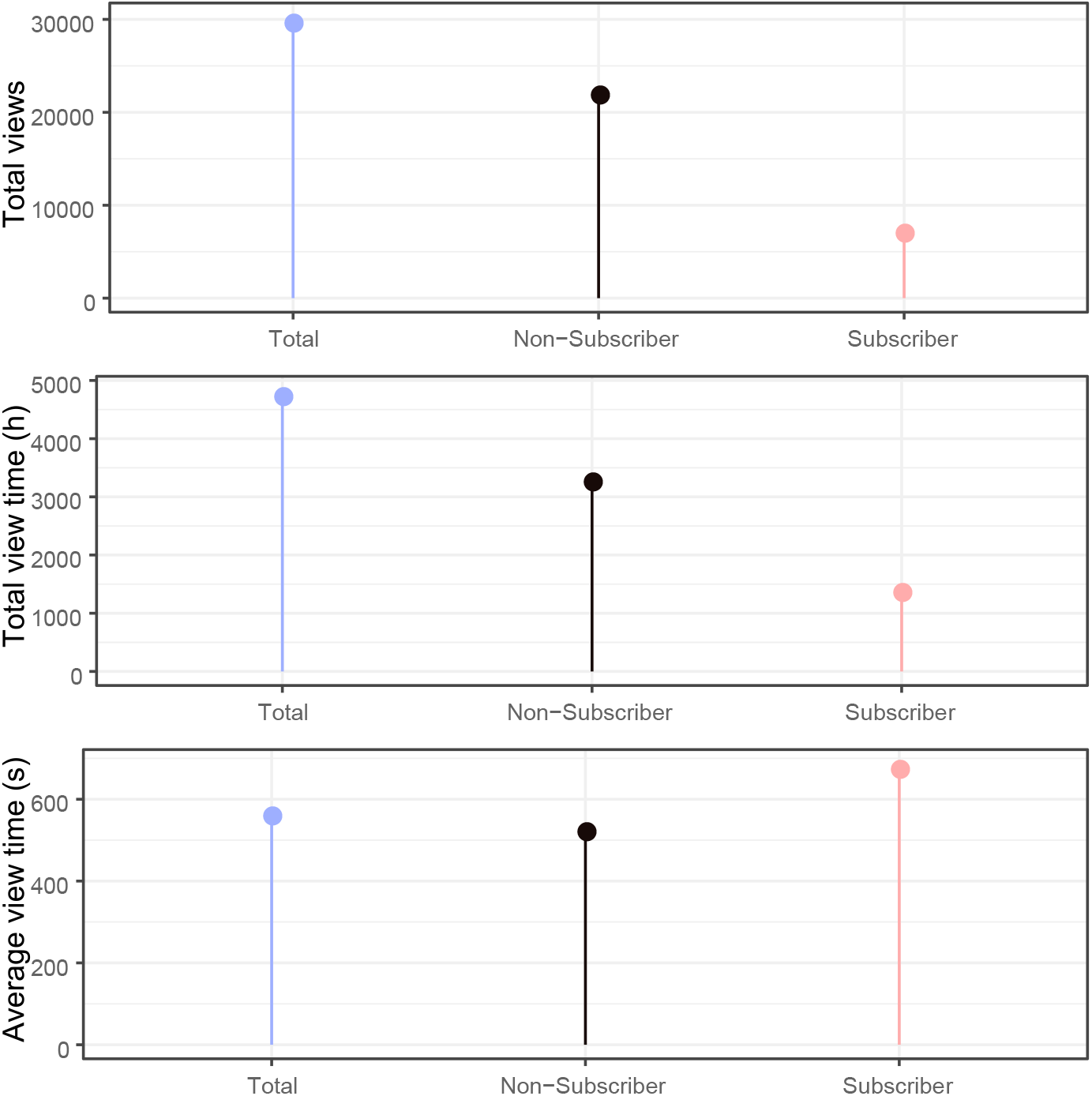
Differences between non-subscribers and subscribers regarding total views, total view time, and average view time.

### 3.3 Data on viewers

Specific viewer data (e.g., location) is only available for viewers with a YouTube account, which does bias our data. WfW episodes were accessed from 11 different countries in Europe, North America, South America, and Oceania (Fig. 6). The majority of views were from Germany (70.2%), followed by Argentina (0.2%), Switzerland (0.2%), and Spain (0.1%). Less than 0.1% of viewers originated from Austria, Mexico, Norway, Australia, the United Kingdom, the Netherlands, and the United States of America. The country of access was not identifiable for the remaining cases and numbers thus do not add up 145 top 100%.

**Figure 6.**
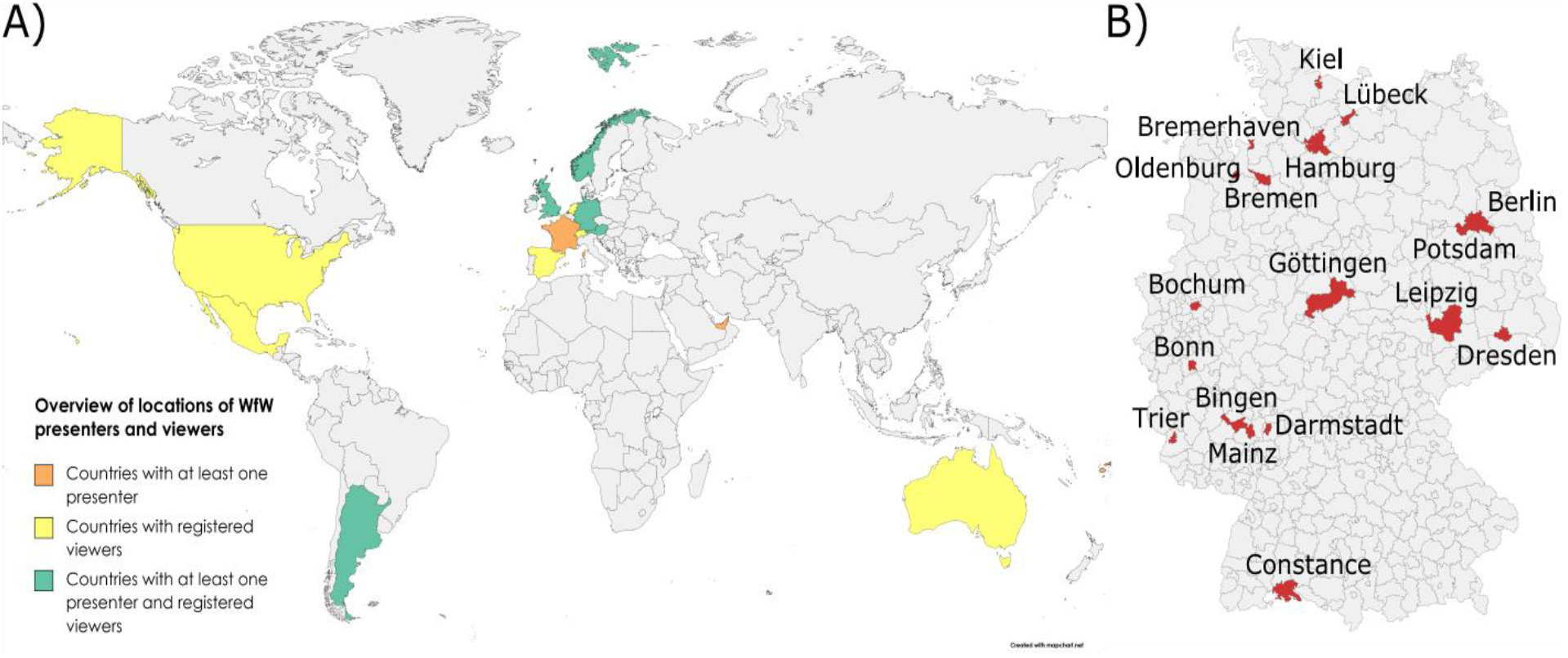
A) World map displaying the geographic location of viewers and presenters. The size of Fiji is enlarged to enhance visibility. B) Map of Germany, with red color displaying the administrative districts of presenters’ affiliations.

The computer is the main medium of accessing WfW with an amount of 16,445 views (54.4%), followed by 10,628 views (35.1%) on smartphones, 1,767 views (5.8%) on tablets, and 1,337 views (4.4%) on TVs. The devices used for the missing 74 views could not be derived. The viewing trend over time is similar for all devices (Fig. 7). Access numbers from the computer and the smartphone rise and fall in similar months. Tablet and TV access is generally much lower, but also more constant. Interestingly, average view times differ strongly between the different devices. The longest average view time is reached by users with TVs (around 17 min), followed by tablets (around 13 min), the computer (around 10 min), and the smartphone (around 6 min).

**Figure 7.**
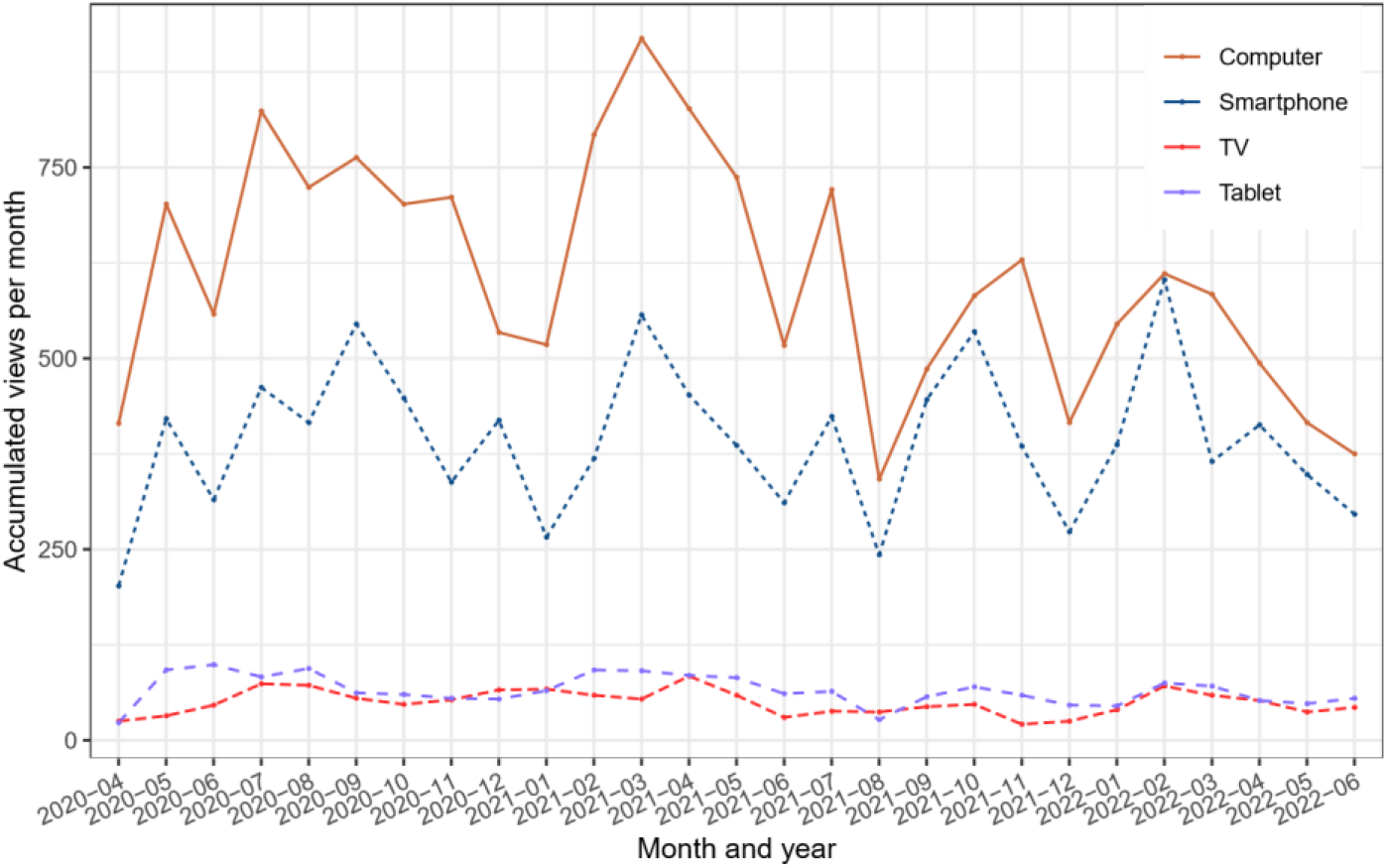
Monthly views classified after the used device. June 2022 only shows data until 09.06.2022.

### 3.4 Topics and presenters

We hosted presentations from different disciplines, with natural sciences as the major topic ranging from biology to glaciology and climate modelling. Other fields included social sciences, psychology, politics, city and landscape planning, aquaculture, and technology development. Several talks were highly interdisciplinary and connected different research fields. These talks were grouped in more than one topic. The main topics were biology and biodiversity, technology and innovation, the polar regions, and climate change impacts with 25, 22, 20, and 12 talks, respectively. The 100 WfW episodes were presented by 91 speakers while 9 scientists presented multiple times. Presenters were mainly senior researchers (46), i.e., > 7 years after obtaining a PhD, ECRs (39), and Non-Academics (15). They represent 38 different affiliations from 26 locations, mainly from Europe (23) and especially Germany (17) (Fig. 6) plus presenters from Argentina, Fiji, and the United Arab Emirates. Twelve presentations were held in English (Table 1). In the first month, presentations were done by the WfW team to kick-start the format; consequentially the focus was on AWI-related research. Over time, covered topics spanned biology, oceanography, glaciology, climate modelling, permafrost, sea ice physics, geology, and chemistry. In addition, two contributions were related to the Multidisciplinary drifting Observatory for the Study of Arctic Climate (MOSAiC) expedition and presented photos, videos and first results (e.g., Nicolaus et al., 2022; Rabe et al., 2022).

We created playlists to group streams by topic. However, the number of videos per playlist varies, and videos can belong to different playlists; making statistics difficult to compare. The playlist with the highest number of videos are “Klimawandel und Gesellschaft” (Climate Change and Society, n=32) and “Arktis und Antarktis” (Arctic and Antarctic, n=26) (Table 1).

## 4 Discussion and Outlook

After two years of livestreaming, WfW has been established as an easily accessible and regular source of up-to-date scientific content. The channel’s success is mirrored by the positive trend of subscribers and the continuous rise in total views (Fig. 2 and 4). Being a non-commercial channel only maintained by voluntary researchers from AWI, WfW does not aim to reach the level of professional, German-speaking channels, such as MaiLab or Quarks. However, the approach of delivering content regularly has resulted in continuous long-term growth (Fig. 2 and 4). WfW has been able to address current topics, such as the federal election in Germany in September 2021^2^, the finding of an extensive colony of fish nests in Antarctica^3^, and the rediscovery of the ship wreck of the polar explorer *Endurance*^4^.

### 4.1 Direct science communication

WfW enables a direct way of science communication. Scientific work is not rewritten by journalists or simply being quoted, but directly and personally presented by researchers. Such direct contact between the public and science is usually limited in popular media and if happening, it is seldom longer than a couple of minutes. Researchers only have limited capacities for voluntary science communication besides their normal tasks; hence, media platforms often inform about science topics without directly consulting a researcher from a specific field. Especially in media, but also in research, some topics attract much public attention, such as the possible crossing of a tipping point of the Greenland Ice Sheet (e.g., King et al., 2020) or the finding of a vast fish breeding colony in Antarctica (Purser et al., 2022). While WfW also presents such science, we similarly broadcast research from niche topics. At WfW, among many more, the following topics were discussed: deformation in ice, blue carbon, sea floor organisms, country-city migration in India, the imperial way of living, and modelling of phytoplankton community changes and species distribution. Most topics are related to climate change and/or biodiversity loss, but are barely discussed in the public media. Providing a platform to present the state-of-the-art research on such topics for the experts from these fields is thus a win-win situation for the audience and the presenter.

### 4.2 Range of topics and presenters

Major perks of WfW are the broad range of themes and methods presented, and the possibility to get to know the people behind the science, i.e. making research more tangible. The topics are rooted within the overall theme of the changing Earth system and habitats connected to climate change and biodiversity crisis, thus allowing the audience to learn about a new field of research each week. Furthermore, it represents many facets of climate-related research and the need for interdisciplinary collaborations to the layman. These different research fields rely on a variety of methods and range from laboratory work, to modelling, and to field work and observational data. WfW communicates different scientific disciplines and highlights the importance of interdisciplinary research incorporating both observational and computed data from diverse disciplines.

### 4.3 Impact

A crucial motivation for science communication is to maximise reach while delivering accurate, easily understandable, and non-populist content. However, what constitutes a significant impact on the general public? Realistic goals are necessary to set the right conditions so that the expectations of the audience and the host are met. A medium such as WfW can not, and tries not, to compete with commercial products or federally funded platforms. The strengths of WfW are a wide variety of topics, interactivity, easy access, and a minimum of obstacles in access and participation. It is furthermore a bottom-up and grass root approach driven by self-motivation. The steady increase in subscriptions and total views demonstrate a successful strategy. Reaching hundreds of people with such a diversity of topics on a weekly basis for over two years is a rewarding endeavour for our group of volunteers while working full-time in academia.

The weekly output, fixed at a specific time and date, is probably one of the main reasons for the success of WfW. Welbourne and Grant (2016) show that videos with consistent science communicators are more popular than videos without a regular communicator. The hosting of WfW by a small group of scientists creates regularity and a sense of familiarity. Nonetheless, having the same moderation team every week is logistically impossible and also not desired, to avoid monotony. Specifically, WfW aims at including a diversity of scientists from different disciplines and career stages.

An active social media presence is one of the most critical points for WfW. Twitter, Instagram, Facebook, and the local media are valuable tools to advertise each episode and the channel. In addition, hosting publicly known researchers showed a positive effect resulting in more than average live views. Such “high impact events” can thus help to gain live views and new subscribers to the channel.

### 4.4 Challenges

We identified three major challenges in establishing and maintaining a successful online outreach medium: 1) keeping a steady and high output, 2) generating new views and subscriptions, 3) balancing seasonal and weather-related ups and downs in views, 4) organising three live hosts and a presenter each Thursday evening, especially upon unforeseen cancellations. Minor challenges are the recruiting of new presenters, the funding of the Zoom account, and the development of a concept to deliver shorter videos.

We have been contacted by companies which aimed to present their products or approaches to solve specific issues. These requests are moral dilemmas, because they are somewhere between research and business. To avoid being hijacked as a marketing platform, such requests are evaluated thoroughly. So far, no purely commercial product has been presented. WfW refuses to advertise for companies that focus on sustainability topics. However, some presenters have informed about their work in new start-ups as part of their research.

### 4.5 Improving public and trans-disciplinary science communication and other skills

In addition to bringing science to the living room at any time and date, WfW is also an education tool for researchers. Presenters enhance their presentation skills, improve their ability to speak in easy language, and expand their own network and communication range. Especially ECRs can sharpen their skills in a moderated and safe environment while choosing the topic and way of presenting themselves. The AWI graduate school *POLMAR* has incorporated WfW in its catalogue allowing doctoral researchers to gain credit points for successful presentations at WfW. Just as with every other skill, training is needed to improve the abilities to become a good science communicator. A platform such as WfW is an easily accessible starting point to improve science communication skills. Furthermore, hosts benefit from establishing new collaborations and scientific exchange. WfW already resulted in one scientific collaboration on the chemical and microbiological coupling between sea and atmosphere in the Arctic. This project has been initiated between the presenter and one of the WfW hosts following episode 51, resulting in joint sampling campaigns on Svalbard subsequently presented in episode 91. Furthermore, in such a large institute as AWI with over 1000 employees, this format has also fostered the interdisciplinary communication and discussions.

### 4.6 Lessons learned and ideas for the future

The success of WfW shows that there is broad public interest in a platform offering a variety of scientific topics. However, expanding and improving the channel while keeping it relevant and practically feasible is a significant challenge, and two approaches are discussed here.

To reduce the regular workload and appeal to the short-term attention span of only a few minutes, which is especially the case for smartphone users, we will try new formats in the channel. One idea is to briefly introduce a researcher and their topic via social media. Interested followers can then ask direct questions to the researcher in an interview, and an edited short version of this interview will be uploaded. This approach maintains the “eye to eye” level of WfW while being much shorter than the usual presentation and discussion videos (5 vs 45 min). Our data shows that the average viewer does not follow the entire video. Even though this depends on the device used for access, shorter content has great potential. Therefore, content will be fewer but more focused on specific sections. Theoretically, this approach is more interactive since followers’ engagement on platforms like Twitter and Instagram, is higher and quicker. Finally, this approach could attract a wider audience while requiring less time and capacity of the WfW team, because the content can be created independently in advance. However, a challenge here is to compete with highly professionalised content already available. The focus would be therefore less on the quality of graphics and design of the videos, but on the authenticity of the presenting scientist and their work.

Self-directed videos of typical lab sessions, measurement procedures, or during expeditions are another promising approach. Creators will record videos using a smartphone and are thus free to choose the focus of their video. The maximum duration is a couple of minutes, which would especially cater for users using smartphones or tablets. As more than one third of the total views origin from smartphones (Fig. 7), content adapted to this medium has therefore considerable potential. The WfW team will edit the videos by adding subtitles, links, and background music to enhance the overall attractiveness, where subtitles would again mainly target smartphone users. These approaches might offer a broader, more diverse catalogue of content to complement the current presentation and discussion videos. However, to maintain the WfW main objective, regular livestreams are still envisioned every second week while question-oriented and self-directed videos take turns in the open slots.

## 5 Conclusions

“Wissenschaft fürs Wohnzimmer” (WfW) is an interactive YouTube channel offering weekly livestreams with a focus on climate and biodiversity. Major aims were to establish a low-threshold, interactive platform for attractive science communication during the first COVID-19 lockdown in early 2020. However, the channel continuous to run successfully and has established itself as a recognised platform. Subscriptions and views are continuously rising since the start of the channel, times of inactivity (e.g., Christmas and summer breaks) are the only time of stagnation. Our success is highlighted by constant numbers of live viewers, and up to several hundred clicks after only a few weeks. Computer and mobile users are the main target audience, even though TV users watch on average twice as long. We identified major lessons learned from two years of WfW and discuss ways to further improve the quality, accessibility, and impact of WfW. This will be helpful for similar media and might allow easier access for novel channels or people new to science communication.

## Data availability

Data will be made publicly available as open access via the Pangaea repository (www.pangaea.de) after acceptance of the manuscript.

## Video supplement

All episodes of WfW are publicly available on *https://www.youtube.com/c/WissenschaftfürsWohnzimmer*

## Author contributions

NS, MW, SJ, FP, CP, JCM, MS, MZ, MK, RM, and BS are the organisers of WfW, designed the study, and collected the data. NS wrote the manuscript with contributions from all co-authors.

## Acknowledgements

We thank all WfW supporters, AWIs4Future, and all guests presenting their research. NS thanks the Helmholtz Junior Research group “The effect of deformation mechanisms for ice sheet dynamics” (VH-NG-802).

## Footnotes

1 https://soundcloud.com/livingroom7a/wfw

2 streamed on 29.07.2021

3 streamed on 03.02.2022

4 streamed on 05.05.2022

## References

• Bickford, D., Posa, M. R. C., Qie, L., Campos-Arceiz, A., and Kudavidanage, E. P.: Science communication for biodiversity conservation, Biological Conservation, 151, 74–76, https://doi.org/10.1016/j.biocon.2011.12.016, 2012.

• Bubela, T., Nisbet, M. C., Borchelt, R., Brunger, F., Critchley, C., Einsiedel, E., Geller, G., Gupta, A., Hampel, J., Hyde-Lay, R., Jandciu, E. W., Jones, S. A., Kolopack, P., Lane, S., Lougheed, T., Nerlich, B., Ogbogu, U., O’Riordan, K., Ouellette, C., Spear, M., Strauss, S., Thavaratnam, T., Willemse, L., and Caulfield, T.: Science communication reconsidered, Nature Biotechnology, 27, 514–518, https://doi.org/10.1038/nbt0609-514, 2009.

• Burgess, J., Green, J., Jenkins, H., and Hartley, J.: YouTube: Online Video and Participatory Culture (DMS - Digital Media and Society), Polity Press, Cambridge, 2009.

• Fischhoff, B.: The sciences of science communication, Proceedings of the National Academy of Sciences of the United States of America, 110, 14033–14039, https://doi.org/10.1073/pnas.1213273110, 2013.

• Fischhoff, B. and Scheufele, D. A.: The science of science communication, Proceedings of the National Academy of Sciences, 110, 14031–14032, https://doi.org/10.1073/pnas.1312080110, 2013.

• Hagedorn, G., Loew, T., Seneviratne, S. I., Lucht, W., Beck, M.-L., Hesse, J., Knutti, R., Quaschning, V., Schleimer, J.-H., Mattauch, L., Breyer, C., Hübener, H., Kirchengast, G., Chodura, A., Clausen, J., Creutzig, F., Darbi, M., Daub, C.-H., Ekardt, F., Göpel, M., Judith N., H., Hertin, J., Hickler, T., Köhncke, A., Köster, S., Krohmer, J., Kromp-Kolb, H., Leinfelder, R., Mederake, L., Neuhaus, M., Rahmstorf, S., Schmidt, C., Schneider, C., Schneider, G., Seppelt, R., Spindler, U., Springmann, M., Staab, K., Stocker, T. F., Steininger, K., Hirschhausen, E. v., Winter, S., Wittau, M., and Zens, J.: The concerns of the young protesters are justified: A statement by Scientists for Future concerning the protests for more climate protection, GAIA - Ecological Perspectives for Science and Society, 28, 79–87, 305 https://doi.org/10.14512/gaia.28.2.3, 2019.

• IPBES: Summary for policymakers of the global assessment report on biodiversity and ecosystem services of the Intergovernmental SciencePolicy Platform on Biodiversity and Ecosystem Services., Tech. rep., IPBES secretariat, Bonn, Germany, https://zenodo.org/record/3553579#.YfmYTerMI2w, 2019.

• IPCC: IPCC 13 Observations: Cryosphere, Climate Change 2013: The Physical Science Basis. Contribution of Working Group I to the Fifth Assessment Report of the Intergovernmental Panel on Climate Change, pp. 317–382, https://doi.org/10.1017/CBO9781107415324.012, 2013.

• IPCC: Climate Change 2021: The Physical Science Basis. Contribution of Working Group I to the Sixth Assessment Report of the Intergovernmental Panel on Climate Change, Tech. rep., Intergovernmental Panel on Climate Change, Cambridge, United Kingdom and New York, NY, USA, https://doi.org/10.1017/9781009157896, 2021.

• King, M. D., Howat, I. M., Candela, S. G., Noh, M. J., Jeong, S., Noël, B. P. Y., van den Broeke, M. R., Wouters, B., and Negrete, A.: Dynamic ice loss from the Greenland Ice Sheet driven by sustained glacier retreat, Communications Earth & Environment, 1, 1–7, https://doi.org/10.1038/s43247-020-0001-2, 2020.

• Ladle, R. J., Jepson, P., and Whittaker, R. J.: Scientists and the media: the struggle for legitimacy in climate change and conservation science, Interdisciplinary Science Reviews, 30, 231–240, https://doi.org/10.1179/030801805X42036, 2005.

• Lahrach, Y. and Furnham, A.: Are modern health worries associated with medical conspiracy theories? Journal of Psychosomatic Research, 99, 89–94, https://doi.org/10.1016/j.jpsychores.2017.06.004, 2017.

• Lenton, T. M., Held, H., Kriegler, E., Hall, J. W., Lucht, W., Rahmstorf, S., and Schellnhuber, H. J.: Tipping elements in the Earth’s climate system, Proceedings of the National Academy of Sciences, 105, 1786–1793, https://doi.org/10.1073/pnas.0705414105, 2008.

• Lenton, T. M., Rockström, J., Gaffney, O., Rahmstorf, S., Richardson, K., Steffen, W., and Schellnhuber, H. J.: Climate tipping points — too risky to bet against, Nature, 575, 592–595, https://doi.org/10.1038/d41586-019-03595-0, 2019.

• Maynard, A. D.: How to Succeed as an Academic on YouTube, Frontiers Communication, 5, 1–9, https://doi.org/10.3389/fcomm.2020.572181, 2021.

• Nicolaus, M., Perovich, D. K., Spreen, G., Granskog, M. A., von Albedyll, L., Angelopoulos, M., Anhaus, P., Arndt, S., Jakob Belter, H., Bessonov, V., Birnbaum, G., Brauchle, J., Calmer, R., Cardellach, E., Cheng, B., Clemens-Sewall, D., Dadic, R., Damm, E., de Boer, G., Demir, O., Dethloff, K., Divine, D. V., Fong, A. A., Fons, S., Frey, M. M., Fuchs, N., Gabarró, C., Gerland, S., Goessling, H. F., Gradinger, R., Haapala, J., Haas, C., Hamilton, J., Hannula, H. R., Hendricks, S., Herber, A., Heuzé, C., Hoppmann, M., Høyland, K. V., Huntemann, M., Hutchings, J. K., Hwang, B., Itkin, P., Jacobi, H. W., Jaggi, M., Jutila, A., Kaleschke, L., Katlein, C., Kolabutin, N., Krampe, D., Kristensen, S. S., Krumpen, T., Kurtz, N., Lampert, A., Lange, B. A., Lei, R., Light, B., Linhardt, F., Liston, G. E., Loose, B., Macfarlane, A. R., Mahmud, M., Matero, I. O., Maus, S., Morgenstern, A., Naderpour, R., Nandan, V., Niubom, A., Oggier, M., Oppelt, N., Pätzold, F., Perron, C., Petrovsky, T., Pirazzini, R., Polashenski, C., Rabe, B., Raphael, I. A., Regnery, J., Rex, M., Ricker, R., Riemann-Campe, K., Rinke, A., Rohde, J., Salganik, E., Scharien, R. K., Schiller, M., Schneebeli, M., Semmling, M., Shimanchuk, E., Shupe, M. D., Smith, M. M., Smolyanitsky, V., Sokolov, V., Stanton, T., Stroeve, J., Thielke, L., Timofeeva, A., Tonboe, R. T., Tavri, A., Tsamados, M., Wagner, D. N., Watkins, D., Webster, M., and Wendisch, M.: Overview of the MOSAiC expedition: Snow and sea ice, Elementa, 10, https://doi.org/10.1525/elementa.2021.000046, 2022.

• Purser, A., Hehemann, L., Boehringer, L., Tippenhauer, S., Wege, M., Bornemann, H., Pineda-Metz, S. E., Flintrop, C. M., Koch, F., Hellmer, H. H., Burkhardt-Holm, P., Janout, M., Werner, E., Glemser, B., Balaguer, J., Rogge, A., Holtappels, M., and Wenzhoefer, F.: A vast icefish breeding colony discovered in the Antarctic, Current Biology, https://doi.org/10.1016/j.cub.2021.12.022, 2022.

• Rabe, B., Heuzé, C., Regnery, J., Aksenov, Y., Allerholt, J., Athanase, M., Bai, Y., Basque, C., Bauch, D., Baumann, T. M., Chen, D., Cole, S. T., Craw, L., Davies, A., Damm, E., Dethloff, K., Divine, D. V., Doglioni, F., Ebert, F., Fang, Y.-C., Fer, I., Fong, A. A., Gradinger, R., Granskog, M. A., Graupner, R., Haas, C., He, H., He, Y., Hoppmann, M., Janout, M., Kadko, D., Kanzow, T., Karam, S., Kawaguchi, Y., Koenig, Z., Kong, B., Krishfield, R. A., Krumpen, T., Kuhlmey, D., Kuznetsov, I., Lan, M., Laukert, G., Lei, R., Li, T., Torres-Valdés, S., Lin, L., Lin, L., Liu, H., Liu, N., Loose, B., Ma, X., McKay, R., Mallet, M., Mallett, R. D. C., Maslowski, W., Mertens, C., Mohrholz, V., Muilwijk, M., Nicolaus, M., O’Brien, J. K., Perovich, D., Ren, J., Rex, M., Ribeiro, N., Rinke, A., Schaffer, J., Schuffenhauer, I., Schulz, K., Shupe, M. D., Shaw, W., Sokolov, V., Sommerfeld, A., Spreen, G., Stanton, T., Stephens, M., Su, J., Sukhikh, N., Sundfjord, A., Thomisch, K., Tippenhauer, S., Toole, J. M., Vredenborg, M., Walter, M., Wang, H., Wang, L., Wang, Y., Wendisch, M., Zhao, J., Zhou, M., and Zhu, J.: Overview of the MOSAiC expedition: Physical oceanography, Elementa: Science of the Anthropocene, 10, 1–31, https://doi.org/10.1525/elementa.2021.00062, 2022.

• Similarweb: Top Website-Ranking, https://www.similarweb.com/de/top-websites/(Dateaccessed:20.06.2022), 2022.

• Uscinski, J. E., Douglas, K., and Lewandowsky, S.: Climate Change Conspiracy Theories, in: Oxford Research Encyclopedia of Climate Science, Oxford University Press, https://doi.org/10.1093/acrefore/9780190228620.013.328, 2017.

• Vaidyanathan, G.: The world’s species are playing musical chairs: how will it end? Nature, 596, 22–25, https://doi.org/10.1038/d41586-02102088-3, 2021.

• Welbourne, D. J. and Grant, W. J.: Science communication on YouTube: Factors that affect channel and video popularity, Public Understanding of Science, 25, 706–718, https://doi.org/10.1177/0963662515572068, 2016.

